# Noninvasive prenatal diagnosis for Haemophilia A by a haplotype-based approach using cell-free fetal DNA in maternal plasma

**DOI:** 10.1101/563429

**Authors:** Chao Chen, Jun Sun, Yun Yang, Lu Jiang, Fengyu Guo, Yaping Zhu, Dan Li, Renhua Wu, Rong Lu, Mei Zhao, Fang Chen, Peixiang Ni, Zhihui He, Zhiyu Peng

**Author notes:** The first three authors contributed equally to this work. **Correspondence:** Zhiyu Peng; Zhihui He.

## Abstract

Because of the background of maternal DNA and technical obstacles in determining the inversion breakpoint in plasma, the direct detection of intron 22 inversion in the *F8* gene is impracticable. A haplotype-based method could potentially solve this problem by analyzing mutation-linked polymorphism sites. This study aimed to demonstrate the feasibility of haplotype-based noninvasive prenatal diagnosis for Haemophilia A. Two families affected by Haemophilia A participated in our study. Maternal haplotypes associated with pathogenic mutations were generated using targeted region genotypes of the mothers and the probands. Combined with the maternal pathogenic haplotypes, a hidden Markov Model was constructed to deduce fetal haplotypes using high-coverage targeted sequencing of maternal plasma DNA. The presence of a pathogenic haplotype in a male indicated an affected fetus, and that in a female indicated a carrier. Prenatal diagnoses were confirmed with amniocentesis and long-distance PCR. The haplotype-based noninvasive prenatal diagnosis of Haemophilia A was successfully performed. One fetus was identified to be Hemophilia A-negative and the other fetus was identified as a carrier; the results were confirmed by amniocentesis. Our research demonstrated the feasibility of noninvasive prenatal diagnosis of Hemophilia A caused by intron 22 inversion in the *F8* gene using a haplotype-based approach.

## 1. Introduction

Haemophilia A (HA) is one of the most common X-linked recessive inherited bleeding disorders, with an incidence of approximately 1/5,000 in males [1], and is caused by a genetic mutation of the *F8* gene (NM_000132.3) that results in deficiency of coagulation factor VIII in plasma. The clinical manifestations include spontaneous or traumatic bleeding in several organs. Prenatal diagnosis is important for HA families because of the lack of an effective treatment. However, an invasive prenatal diagnosis approach such as chorionic villus sampling (CVS) and amniocentesis (AF) has a small risk of miscarriage and infection [2]. The discovery of cell-free fetal DNA (cff-DNA) in maternal plasma makes noninvasive prenatal diagnosis (NIPD) possible [3]. The intron 22 inversion in the *F8* gene is found in approximately 45% of severe HA patients [4]. Because of the background of maternal DNA [5] and technical obstacles in determining the inversion breakpoint in plasma, the direct detection of HA is impracticable. A haplotype-based method could potentially solve this problem by analyzing mutation-linked polymorphism sites [6]. This study describes the few uses of NIPD for HA by a haplotype-based approach and demonstrates that our approach is an affordable strategy for the clinically feasible NIPD of X-linked disorders.

## 2. Results

### 2.1. Sequencing data

For maternal plasma, we obtained a mean 41.35 million data mapped to the target region with a mean depth of 376× and 99.48% coverage with more than 20 reads. A mean depth of 151× and 97.93%with 20× coverage was obtained in genomic DNA of pregnant women, their brothers and their spouses (Table 1). The data were enough for subsequent analysis.

**Table 1.** Sequencing data of studied families

| Family | Sample | Target region data (M) | Mean depth (x) | 20× Coverage fraction | Mean depth of chr Y | 20× Coverage fraction of chr Y | SNPs | GW | Fetal DNA fraction |
| --- | --- | --- | --- | --- | --- | --- | --- | --- | --- |
| F01 | mother | 21.29 | 193 | 98.70% | 0 | 0 | 248 | 17+2 | 11.09% |
|  | father | 15.50 | 141 | 98.31% | 45 | 72.2% |  |  |  |
|  | proband | 15.14 | 138 | 98.11% | 43 | 71.1% |  |  |  |
|  | plasma | 29.93 | 272 | 99.37% | 27 | 42.59% |  |  |  |
|  | AF | 11.00 | 100 | 96.80% | 114 | 96.74% |  |  |  |
| F02 | mother | 22.73 | 206 | 98.79% | 0 | 0 | 106 | 17+3 | 9.37% |
|  | father | 10.22 | 93 | 95.81% | 102 | 95.4% |  |  |  |
|  | proband | 14.91 | 135 | 97.83% | 153 | 97.4% |  |  |  |
|  | plasma | 52.77 | 479 | 99.59% | 0 | 0 |  |  |  |
|  | AF | 21.06 | 191 | 98.63% | 0 | 0 |  |  |  |

Notes: AF, Amniotic fluid; GW, gestational weeks; the proband was the brother of each pregnant woman; SNPs represents the number of SNPs detected in the studied families.

### 2.2 Determination of fetal sex and DNA fraction

The mean depth and 20× coverage mapped to chromosome Y in F01 plasma was 27× and 42.59%. The fetus of F01 was inferred to be male. However, no reads were mapped to the target region of chromosome Y, so the fetus of F02 was determined to be female. The fetal DNA fractions of F01 and F02 were 11.09% and 9.37%, respectively.

### 2.3 Haplotype-based noninvasive prenatal diagnosis

The maternal haplotype could be constructed based on these SNPs. The number of informative phased SNPs used for NIPD for F01 and F02 was 144 and 49, respectively. In F01, the number of SNPs (Table 2, Figure 1) indicating that the fetus inherited the maternal wild-type haplotype (Hap1) was 144, while none supported the pathogenic haplotype (Hap0). The fetus could be subsequently predicted to be unaffected with HA considering the fetus’s male sex. In F02, the number of SNPs (Table 2, Figure 1) indicating that the fetus inherited the maternal pathogenic haplotype (Hap0) was 49, while none supported the wild-type haplotype (Hap1). The fetus could be subsequently predicted to be a carrier of HA considering the fetus’s female sex.

**Table 2.** The NIPD results from *F8* gene analysis

| Family | Fetal Sex | Phased SNPs<sup>a</sup> | SNPs For Hap0<sup>b</sup> | SNPs For Hap1<sup>c</sup> | NIPD results | Predicted results |
| --- | --- | --- | --- | --- | --- | --- |
| F01 | male | 144 | 0 | 144 | Hap1 | normal |
| F02 | female | 49 | 49 | 0 | Hap0 | carrier |

Notes: a represents the number of SNPs that were used to predict parental haplotypes; b represents the number of SNPs that supported the pathogenic haplotype (Hap0); c represents the number of SNPs that SNPs that supported the normal haplotype (Hap1).

**Figure 1:**
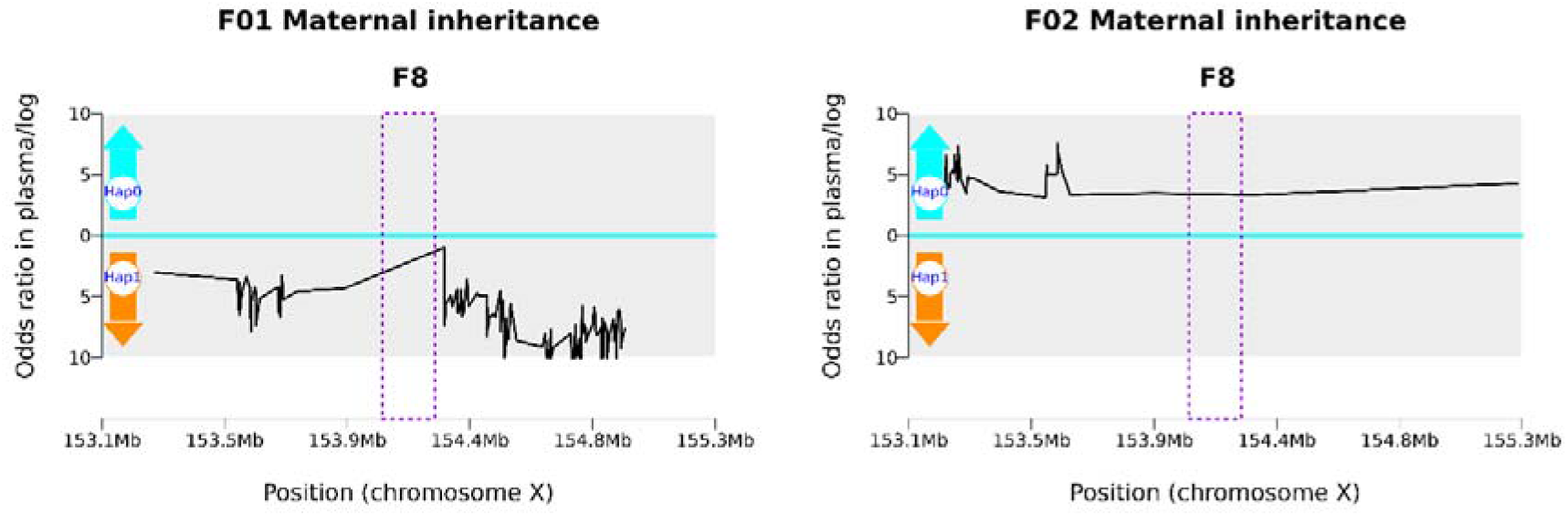
Results of haplotype-based NIPD of Haemophilia A. The X-axis represents the target region of our customized probe, and blue vertical lines represent the interval of the *F8* gene; the Y-axis represents the logarithm of the ratios of fetal-inherited maternal haplotype. Hap0 represents the pathogenic haplotype, and Hap1 represents the wild-type haplotype. The black lines above zero indicate that the fetus inherited the pathogenic haplotype (Hap0), and the black lines below zero indicate that the fetus inherited the normal haplotype (Hap1).

### 2.4 Invasive prenatal diagnosis

Prenatal diagnosis performed by LD-PCR of F01 was indicated to be a fetus unaffected by HA (Figure 2A). The fetus of F02 was diagnosed as a carrier of HA (Figure 2B).

**Figure 2:**
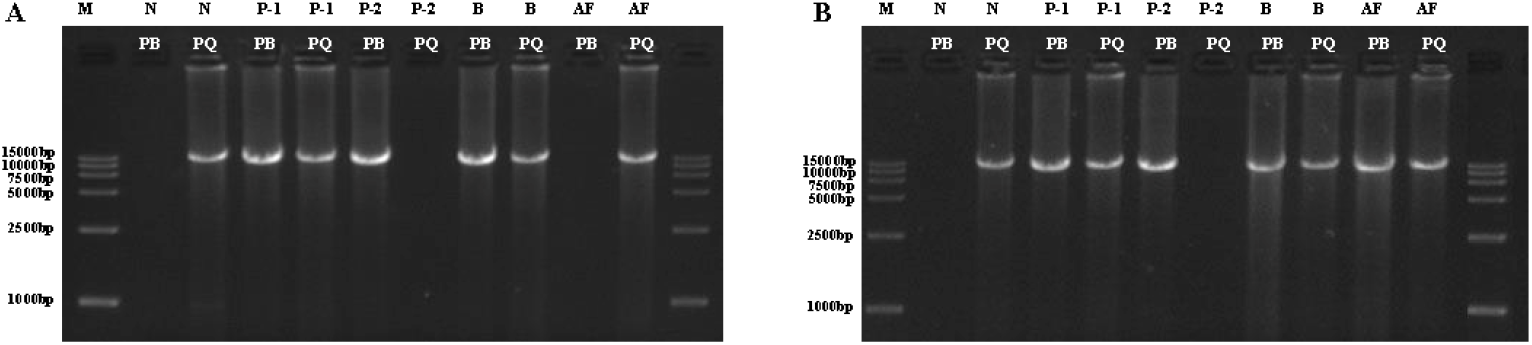
Results of invasive diagnosis of studied families. PB was amplified by primers P and B, and PQ was amplified by primers P and Q. M: DL15000 Marker; Lane N: normal control; Lane P-1: female carrier control; Lane P-2, male patient control; Lane B, Mother; Lane AF: amniotic fluid. (A) Electrophoresis results of LD-PCR of amniotic fluid (F01). (B) Electrophoresis results of LD-PCR of amniotic fluid (F02).

## 3. Discussion

The discovery of cell-free fetal DNA (cff-DNA) in maternal plasma bring a new resource for noninvasive prenatal diagnosis. Heretofore, several studies have reported the high accuracy of noninvasive prenatal detection of fetal chromosome aneuploidy [7, 8]. In addition, whole-genome sequencing (WGS) of maternal plasma DNA can predict the fetal-specific allele with accuracy of more than 95%, thus providing the possibility of single gene detection in maternal plasma[9]. Nevertheless, the extremely high cost of whole-genome deep sequencing has limited its clinical application. Deep sequencing of target regions is a noninvasive alternative to single-gene-based disease detection, which could enable deep sequencing of interest regions at a lower cost [10]. Lam et al. developed an algorithm for noninvasive prenatal fetal diagnosis of single-gene diseases using target region capture sequencing for single-base variations and indels [11]. Using this method, two fetuses were correctly diagnosed as β-thalassemia carriers.

Cell-free fetal DNA analysis in maternal plasma provides a noninvasive method for assessing fetal sex in HA-associated pregnancies, whereas without mutation-specific confirmatory analysis, the disease status of male fetuses remains unknown. Tsui et al. used microfluidics digital PCR method to perform noninvasive prenatal diagnosis of HA [12]. However, it can only detect point mutations. The high incidence of the *F8* inversion 22 (Inv 22) mutation cannot be detected by this method. Microfluidics digital PCR is expensive as a microfluid digital PCR chip costs thousands of dollars. Our team developed a haplotype-based strategy for one-step prediction of fetal genotype and haplotype based on both parentally heterozygous sites in the maternal plasma [6].Using this method, we successfully predicted the fetal mutation status of many autosomal recessive genetic diseases using maternal plasma, including congenital adrenal hyperplasia, maple syrup urine disease, congenital deafness and Duchenne muscular dystrophy [13–15]. Our haplotype-assisted strategy provides an opportunity for the diagnosis of HA by NIPD. As a simple procedure, samples for the NIPD method are more accessible than those for invasive prenatal diagnosis. NIPD for HA would allow testing in the early weeks of gestation, giving more opportunity and time for making decisions regarding terminating the pregnancy. Moreover, it provides an option for women who refuse invasive methods to have more information about the possibility of an affected fetus.

At present, fetal gene sequencing analysis based on capture technologies are expensive due to the large target region. Hudecova et al. reported a haplotype-based approach combined with ddPCR for NIPD of HA using a 27.3M custom probe set [16]. A small target region of 110.08 kb was designed including the F8 gene coding area, 445 highly heterozygous SNPs distributed within 1 Mb flanking regions of *F8,* and a 10.37 kb chromosome Y-specific region in this research. Therefore, we can reduce the cost of sequencing substantially. Several studies reported NIPD methods for monogenic disorders using direct haplotype phasing based on linked-reads sequencing and targeted locus amplification (TLA)[17–19]. The linked-reads sequencing was too expensive to obstruct clinical transformation. The TLA technology needs customized target kit for different monogenic disorders, which was inconvenient. The intron 22 inversion in the F8 gene is found in approximately 45% of severe HA patients [4]. Technical obstacles in determining the inversion breakpoint using next generation sequencing makes impossible for NIPD of HA using direct haplotype phasing methods. Our research developed the haplotype phasing method based on the trio’s members without direct detection of inversion mutation. All types of mutation including copy number variation, indel inversion and translocation could be detectable using our method.

Sex determination is very important for NIPD of X-linked inherited diseases. We use the specific Y chromosome probe to easily and accurately perform sex determination. The sequencing depth of target region on the Y chromosome are markedly different between male and female fetuses according to cff-DNA sequencing data analysis. It is approximately 0 for female fetus. NIPD may predict fetal sex accurately.

Even though NIPD of HA has many advantages compared with invasive methods, the accuracy needs to be further improved before its clinical application. As accuracy increases and costs reduce, we hypothesize that, in the future, NIPD will be used as a relatively fast and easy alternative for pregnant women who are likely to have fetuses with HA and other X-linked diseases.

## 4. Materials and methods

### 4.1. HA patient recruitment and clinical information

Two families (F01-F02) with individuals diagnosed with HA, including two pregnant women, their affected brothers (proband) and their spouses were recruited. According to the residual plasma FVIII coagulant activity (FVIII: C) [20], the proband of F01 and F02 was classified as moderate (4%) and severe (0.8%), respectively. The causative mutation in the brothers and the pregnant women was previously identified to be an intron 22 inversion in the *F8* gene using targeted sequencing combined with long-distance PCR (LD-PCR). Genetic counseling was given to the families, and the parents decided to take our noninvasive prenatal test. All participants gave their signed informed consent. The ethics committee of The First Affiliated Hospital of Guangzhou Medical University approved this study. The 5 mL blood samples obtained from each family member, including mother, father and proband, were transported to BGI-Tianjin. Maternal plasma was separated using a double-centrifugation method and stored frozen. Amniotic fluid was obtained through amniocentesis for routine prenatal diagnosis. Invasive prenatal diagnosis and NIPD were performed in a double-blind manner by two independent investigators.

### 4.2. Library preparation and targeted sequencing

In this study, we used haplotype-based noninvasive prenatal testing of HA using a 110.08 kb SeqCap Kit (Roche, Basel, Switzerland) for the *F8* gene coding region and 445 surrounding highly heterozygous SNPs (minor allele frequency, MAF=0.3-0.5) distributed within a 1 M bp region of the X chromosome (Figure 3). Inclusion of a 10.37 kb chromosome Y-specific region facilitated the simultaneous determination of fetal sex.

**Figure 1.**
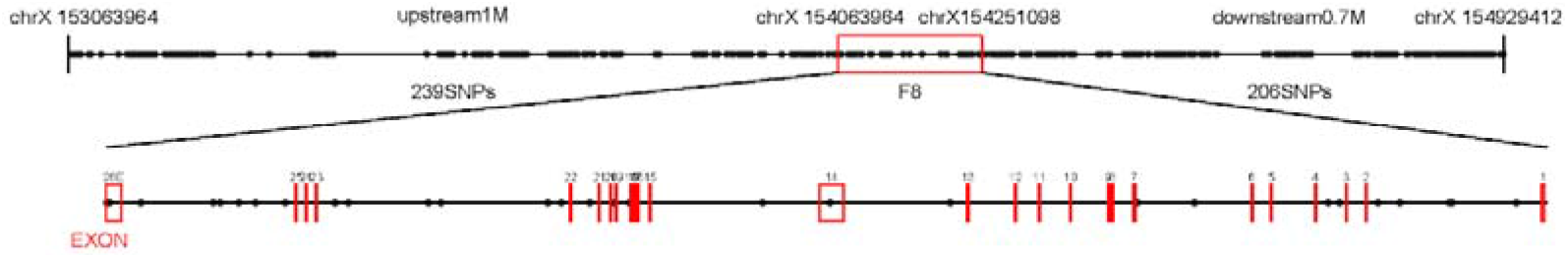
Target region of the *F8* gene and SNPs used for haplotyping. Custom NimbleGen probes were designed that targeted within the 110.08 kb region including the *F8* gene coding region and 445 surrounding highly heterozygous SNPs distributed within a 1 Mb region.

Genomic DNA (gDNA) was extracted from blood or umbilical cord blood using a QIAamp DNA Mini Kit (Qiagen, Dusseldorf, Germany). Cell-free DNA was extracted from maternal plasma with QIAamp Circulating Nucleic Acid Kit (Qiagen, Dusseldorf, Germany). The cell-free DNA library was prepared using the KAPA Library Preparation Kit (Kapa Biosystems, Wilmington, USA). The gDNA library was constructed according to our established method [15]. The libraries of gDNA and cell-free DNA were then pooled together, and then captured using our customized probes. Hiseq2500 platform was used for sequencing. Using BWA software (http://bio-bwa.sourceforge.net/), the sequencing reads were aligned to the human reference genome (hg19). SNP calling was performed using GATK software (https://software.broadinstitute.org/gatk/).

### 4.3. Determination of fetal sex and DNA fractions

The mean depth of the Y chromosome target region was utilized to determine the fetal sex. If the fetus was female, the depth of the Y chromosome region would be extremely low, resulting from short nonspecific region reads that could be mapped to Y chromosome-specific region [21].

The fetal DNA concentration in the maternal plasma fraction was calculated by using SNP sets with different parental homozygosity and the formula ε=2d_F_/(d_F_+ d_M_) [9], where ε represents fetal DNA fraction in plasma, d_F_ is the depth of reads supporting fetal-specific alleles and d_M_ is the depth of reads supporting maternal shared alleles in plasma. Assuming the maternal genotype is AA and the paternal genotype is TT, the fetal genotype should be AT, as the fetal-specific allele was T and the maternal shared allele was A.

### 4.4. Haplotype-based noninvasive prenatal diagnosis of the F8 gene

A noninvasive haplotype-based linkage analysis method as previous described [22] was used to determine fetal haplotype using information regarding SNPs from 1 Mb regions flanking the coding regions of the *F8* gene from probands and parents. We defined haplotype 0 (Hap0) as the pathogenic haplotype and haplotype 1 (Hap1) as the wildtype haplotype. Because candidate Hap0 was inherited from maternal chromosome X, we directly phased the maternal haplotypes of the *F8* gene. Based on maternal haplotype and SNP representation in the plasma, a hidden Markov Model was constructed. We chose the SNP sets that were heterozygous in the mother but homozygous in the father to determine the maternal inheritance. The fetal DNA fraction and sequencing error rate were utilized to determine the maternal haplotype inheritance patterns in the fetus. The probability of each state of maternal inheritance was calculated for each SNP marker in the maternal plasma. The fetal haplotype was deduced using Viterbi algorithm to identify the most hidden states.

### 4.5. Prenatal diagnosis of HA using amniotic fluid

LD-PCR of amniotic fluid using primers P, Q and B [23] was utilized to assess the accuracy of haplotype-based NIPD of HA. Electrophoresis showed a 12 kb band (Fragment PQ) for the wild-type, an 11 kb band (Fragment PB) for male patients and PQ and PB bands for female carriers.

## Author contributions

Conceptualization, Z.P. and Z.H.; Sample collection, D.L., R. L. and M.Z.; Performed the experiments, L.J.; R.W. and C.C.; Analyzed and interpreted the sequence data, F.G.; Y.Z. and P.N. Writing—original draft preparation, C. C., J. S. and Y.Y.; Funding acquisition, Z.H., F.C. and Y.Y.

## Funding

This research was funded by grants from the Natural Science Foundation of Guangdong Province, China (2015A030313472) and Major Technical Innovation Project of Hubei Province (2017ACA097).

## Acknowledgments

We are grateful to the participants and their families. We also thank our coinvestigators for their assistance in research designand data analysis.

## Conflicts of Interest

The authors declare no conflict of interest.

## Notes

#### Summary of Updates

author and communication author updated; Abstract updated;Figure 1,2,3 revised.

